# Context-dependent tradeoffs between morphological plasticity and thermal stress resistance in a reef-building coral

**DOI:** 10.1101/2025.10.07.680072

**Authors:** Jenna Dilworth, Maya Gomez, Ian Combs, Iliyan Hariyani, Joseph Kuehl, Sophia Lee, Tatianna Velicer, Natalie Villafranca, Hanna R. Koch, Erinn M. Muller, Carly D. Kenkel

**Author notes:** corresponding author:, Address: 3616 Trousdale Pkwy, AHF 231, Los Angeles, CA 90089, USA.

## Abstract

In sessile, long-lived organisms, such as reef-building corals, phenotypic plasticity may play a key role in mitigating the impacts of climate change. Plasticity is often context-dependent and has costs and limits, ultimately resulting in adaptive, neutral, or maladaptive effects. However, our understanding of the mechanisms that maintain variation in, and therefore drive the evolution of, plasticity is still limited. We use a field-based transplant experiment to show context-dependent tradeoffs to morphological plasticity in *Acropora cervicornis* under real-world conditions driven by climate change. We find that while morphological plasticity is neutral under some conditions, it trades off with thermal tolerance during moderate temperature stress during a marine heatwave. Additionally, we find that plasticity changes over time, underlining the importance of its temporal dynamics in long-lived organisms. Our results demonstrate a key role for environmental change in the eco-evolutionary dynamics of plasticity by demonstrating context-dependent costs under real-world thermal stress.

## Introduction

As climate change rapidly shifts environmental conditions, organisms must adapt to these changes or relocate to more favorable environments (Parmesan, 2006; Parmesan and Yohe, 2003). In sessile, long-lived organisms, such as reef-building corals, phenotypic plasticity is hypothesized to play an important adaptive role in mitigating climate risk (Chevin et al., 2010; Kelly, 2019). Organisms may respond to environmental change by adjusting their behavior (Gross et al., 2010), physiology (Seebacher et al., 2015), or morphology (Mizutani and Kanaoka, 2018), allowing for phenotypic changes within their lifetime without genetic adaptation (Scheiner, 1993). For example, a population of great tits in Britain advanced its mean egg-laying time via adaptive plasticity to track climate-driven changes in spring temperature (Charmantier et al., 2008). However, our understanding of the mechanisms constraining the evolution of plasticity is still limited. Plastic responses may be adaptive, neutral, or maladaptive (Ghalambor et al., 2007), with ultimate effects on fitness being highly context-dependent (Hendry, 2016). For instance, a study exposing two species of *Lobelia* wildflowers to wet or dry conditions found varying levels of plasticity in several photosynthetic traits, with neutral, adaptive, or maladaptive effects depending on the specific environmental conditions (Caruso et al., 2006). In addition, plasticity is generally associated with costs and limits (DeWitt et al., 1998), but our ability to capture these constraints may be influenced by how we measure plasticity and when we determine its fitness effects (Auld et al., 2009). As climate change continues to accelerate, it will be crucial to understand these dynamics and how the costs or limits associated with plasticity may constrain organisms’ ability to mitigate climate impacts.

The Caribbean branching coral *Acropora cervicornis* is an excellent model to study phenotypic plasticity in the field. Asexual reproduction via clonal fragmentation allows for the production of replicate ramets of the same genotype (Highsmith, 1982; Tunnicliffe, 1981). These ramets are common-gardened before transplantation to natural reefs as part of reef restoration practices (Young et al., 2012), providing the unique opportunity to study plastic responses of the same animal genotype in multiple natural environments. Previous research provided evidence for an adaptive role of plasticity in this system in the Lower Florida Keys by demonstrating that different genotypes of *A. cervicornis* expressed varying levels of morphological plasticity across different environments, and that this plasticity was positively correlated with growth and survival (Million et al., 2022). While this prior study did not find any tradeoffs to plasticity, its costs can be difficult to detect (Auld et al., 2009; Van Buskirk and Steiner, 2009). In animals, costs of plasticity are often highest in stressful environments, likely due to increased constraints in the context of competition or resource limitation (Van Buskirk and Steiner, 2009), suggesting that the costs of morphological plasticity in *A. cervicornis* may be context-dependent and could be revealed under stress. Potential costs under environmental stress are ecologically relevant in reef-building corals, as their obligate endosymbiotic relationship with dinoflagellate algae in the family Symbiodiniaceae (LaJeunesse et al., 2018) can be disrupted at just 1-2°C above local summer maxima in a process known as bleaching (Glynn, 1996). Rising sea surface temperatures are increasing the frequency and severity of bleaching events (Hughes et al., 2018), often leading to mortality or negative fitness impacts (Baird and Marshall, 2002; Leinbach et al., 2021). The continued impacts of mass bleaching events are of particular concern for *A. cervicornis* in Florida, which is now functionally extinct from the state’s reefs (Manzello et al., 2025), with a substantially reduced wild population in which many genets only survive due to restoration programs maintaining them in captivity (Muller et al., 2025).

We conducted a field-based transplant experiment to investigate the context-dependent costs of morphological plasticity in *A. cervicornis* during a real-world thermal stress event. By applying a multivariate method to quantify morphological plasticity at several timepoints (Leung et al., 2020), we show that the relative plasticity expressed by different genotypes at our outplant sites is dynamic over time. Additionally, we demonstrate that the ultimate effects of this plasticity are context-dependent, with a tradeoff between morphological plasticity and resistance to thermal stress evident at moderate levels of temperature stress. These results illustrate how changing environmental conditions may influence the eco-evolutionary dynamics of plasticity in this endangered reef-building coral.

## Materials and Methods

### Outplanting and Monitoring

In October 2022, 20 ramets of consistent size (mean length = 8.3cm, sd = 1.1cm) and without secondary branches (Fig. 1b) of each of ten genotypes of *A. cervicornis* were sourced from Mote Marine Laboratory’s Looe Key *in situ* nursery (24.56257, -81.40009) (Fig. 1a), where they had been common-gardened for 9+ years. The Floridian population of *A. cervicornis* is genetically well-mixed and the genotypes included in this study are reflective of population-level genotypic diversity (Fig. S1). Ramets were photographed with Olympus Tough cameras in batches of ten on racks with scaling bars (Gomez and Kenkel, 2025) adapted from (Million et al., 2021) (Fig. 1b). On October 14th and 15th, ten ramets per genotype were randomly outplanted to one of ten arrays at one of two reef sites in the Lower Florida Keys (n = 100 per site, total n = 200): Dave’s Ledge (24.54672, -81.40159) or Looe Key (24.53154, -81.48356) (Fig. 1a). These sites had demonstrated intermediate levels of outplant survivorship in a previous outplant experiment (Million et al., 2022). PAR measurements recorded with a LICOR sensor (Onset Computer Corporation, Bourne, MA) at outplant depth (18-25 ft) at each site did not overlap with light levels at the nursery (Fig. S2). Masonry nails were hammered into the substrate and corals were affixed to the nails with zip ties. HOBO temperature loggers (Onset Computer Corporation, Bourne, MA) were deployed at both sites to log hourly temperatures throughout the experiment. Both sites were resurveyed in January, April, and June 2023 over 1-3 observation days per visit to re-photograph each outplant with individual scale bars following (Gomez and Kenkel, 2025) (adapted from (Million et al., 2021). Survival was recorded during these surveys and later confirmed with photographs. Outplants where the entire fragment and zip tie and/or masonry nail were missing in January (Dave’s Ledge: n = 15, Looe Key: n = 26; n = 41 total) were considered lost due to technical failure rather than biological mortality and excluded from further analyses.

**Figure 1:**
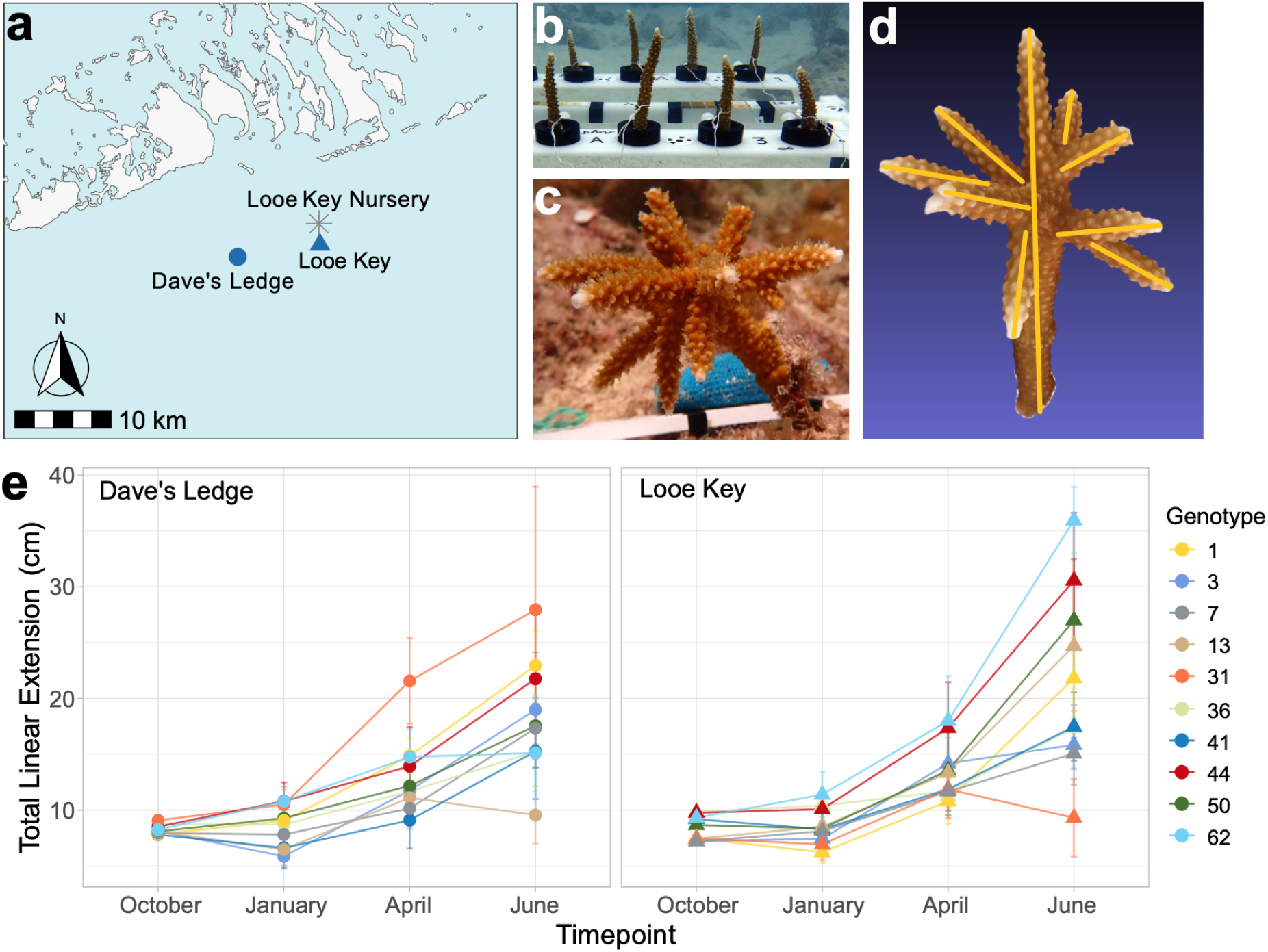
Growth and absolute size vary by outplant site and genotype. **a:** Location of source nursery (grey star) and outplant sites (blue shapes) in the Lower Florida Keys. **b-c:** representative images of experimental replicates in October 2022, before outplanting (**b**) and in June 2023, after 9 months of growth in the field (**c**). **d:** Example 3D model used for phenotyping: yellow lines on each branch represent measurement method for total linear linear extension (TLE). **e:** Genotype mean TLE in cm at each outplant site from initial (October 2022, n = 159) through the following three-month monitoring intervals (January 2023, n = 144; April 2023, n = 141; June 2023, n = 139). Point colors represent genotype, shapes represent site. Error bars show standard error.

### 3D Model Construction and Phenotyping

Photos were sorted into separate photosets for each rack of corals (October 2022) or individual outplant (all following timepoints). Three-dimensional models were generated from photosets using Agisoft Metashape v1.8.3 (Agisoft LLC, St. Petersburg, Russia) (Fig. 1d). Custom Python scripts adapted from (Million et al., 2021) with modifications for automatic scaling with Metashape targets were used to batch process photos using the High Performance Computing resources at the University of Southern California’s Center for Advanced Research Computing. High accuracy was used for point cloud alignment parameters and high quality was selected for depth maps generation. Individual models were visually inspected and rebuilt if there were any in the focal area, scale bar error was >0.01, or model reprojection error was >5 pix. Models were imported into MeshLab (Cignoni et al., 2008) to measure total linear extension (TLE), volume (Vol), and surface area (SA) as described in (Million et al., 2021) (Fig. 1d). For measurements of corals photographed using racks (October 2022), the height of the holder used to attach corals to the rack (Fig. 1b) was added to TLE measurements. Additionally, the height of the holder and diameter of the coral base were used to calculate the volume and curved surface area of a cylinder, which were added to model Vol and SA to account for coral tissue obscured by this attachment method. Breakage type was determined by comparing models to the previous timepoint and categorized by branch ordination as well as branch number, counting from the base of the coral upwards: breakage on the primary branch (P), secondary branches (S), tertiary branches (T), or catastrophic breakage (C, loss of most secondary/tertiary branches and a significant portion of the primary branch). Growth and breakage were calculated as the total gain (growth) or loss (breakage) in total linear extension between two timepoints. Breakage severity was categorized on a scale from 0 (no breakage) to 5 (catastrophic breakage) by integrating breakage type with net growth/breakage data as outlined in Table S4.

### Assessment of Thermal Tolerance

On June 29th 2023, RAW images with Coral Health Chart (CoralWatch, St Lucia, Australia) color scales were taken of each outplant using Olympus Tough cameras to serve as baseline color references in anticipation of a potential bleaching event. After receiving reports of bleaching near our experiment sites, we conducted a non-quarterly monitoring survey of all outplants on July 26th and 28th to assess their response to thermal stress. Mortality was recorded and RAW images with Coral Health Chart color scales were taken of each outplant. Additionally, ∼2 cm tissue samples were taken from branch tips of all living outplants using bone cutters and snap frozen in liquid nitrogen. The amount of temperature stress accumulated at each outplant site was calculated using average daily temperatures calculated from in-water hourly temperatures recorded by HOBO loggers via the experimental Degree Heating Weeks method (Leggat et al., 2022).

To assess visual bleaching severity, outplant color scores were determined in ImageJ (Schneider et al., 2012) using the baseline photos taken in June and marine heatwave photos from July. The Coral Health Chart D1-D6 color reference in each outplant image was used to generate a standard curve of mean grayscale values (Siebeck et al., 2006). Grayscale values of three unshaded areas of each outplant were then measured and converted into a color score corresponding to the Coral Health Chart (Zhang et al., 2022). To assess symbiont to host (S:H) ratios in collected tissue samples, total DNA was extracted using a Qiagen DNeasy PowerBiofilm Kit (Qiagen, Aarhus, Denmark), and relative abundances of *Symbiodinium* and coral host DNA were quantified via qPCR using actin (Winter, 2017) and calmodulin (Palacio-Castro et al., 2021) assays respectively. On a random subset of samples, additional multiplexed *Cladocopium* and *Durusdinium* actin assays (Cunning and Baker, 2020) were run to determine whether there were significant levels of these symbiont genera present. All assays were run for 40 cycles on an Agilent AriaMX (Agilent, Santa Clara, CA, USA) system with two technical replicates and reference dye corrections. C_T_ values were corrected for fluorescence and gene copy number following (Cunning and Baker, 2013).

Though demonstrating similar overall trends, S:H ratios as measured via qPCR and color scores determined from the Coral Health Chart did not correlate (Pearson’s r, p = 0.37). This may be due to the fact that *A. cervicornis* can slough host tissue in addition to the loss of algal symbionts during severe bleaching stress, leading to the loss of both host and symbiont DNA (DeMerlis et al., 2022). Thus, we opted to use color scores for further analyses, since they were determined using the average of three different measurement areas, while tissue samples for S:H ratios were collected from only one area near branch tips, which frequently appear white even in the absence of thermal stress due to lower concentrations of algal symbionts. To integrate mortality and bleaching severity at each outplant site, we used a modified version of the Bleaching Stress Index (BSI) developed by (Humanes et al., 2024). Color scores were binned into the six color categories (c1 - c6) from the Coral Health Chart, while outplants that died from heat stress were given a score of zero (c0). BSI was then calculated for each genotype at each site:

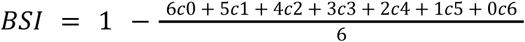

Where c0 to c6 are the proportion of ramets of each genotype that died between June and July or fell into one of the six color score categories from the Coral Health Chart, yielding a BSI ranging from 0 (all ramets dead) to 1 (all ramets alive and visually healthy).

### Replication Statement

| Scale of inference | Scale at which the factor of interest is applied | Number of replicates at the appropriate scale |
| --- | --- | --- |
| Florida population of *Acropora cervicornis* | 10 genotypes at 2 outplant sites | 10 replicate ramets per genotype per site; n = 200 |

### Statistical Analyses

A binomial logistic regression was used to test for significant differences in attachment failure by genotype, outplant site or their interaction. Two-way ANOVAs were used to test for initial differences in TLE, SA, Vol, and SA:Vol by genotype and outplant site. As all traits demonstrated significant initial differences by genotype (Table S1), initial size was included in all subsequent models. Linear mixed-effects models (Bates et al., 2015) were used to investigate the fixed effects of initial TLE as well as timepoint, genotype, site, and their interactions on growth, TLE, SA, Vol, and SA:Vol, with a random intercept for fragment number to account for repeated measures. Linear mixed-effects models were also used to determine the fixed effects of initial size, genotype, site, and the genotype x site interaction on absolute size at the final measurement timepoint in June, with random intercepts by outplant array.

To integrate all morphological traits, we used redundancy analysis (RDA) to assess the effects of genotype and site on multivariate morphology over time. The effect of genotype, site, and their interaction, conditioned on timepoint and initial TLE, on a scaled matrix of all morphological traits (TLE, V, SA, SA:Vol, growth, breakage, and breakage severity) was modeled using the vegan package (Oksanen et al., 2025). Initial TLE was chosen as the conditioning factor to account for baseline differences in morphology because it correlates strongly with other morphological traits in small, unbranched fragments (Million et al., 2021). As the first two RDA axes explained a significant amount of the constrained variation accounted for by genotype, site, and their interaction, genotype-specific plasticity at each timepoint was calculated as the Euclidean distance between the site centroids for each genotype in RDA space following (Leung et al., 2020) (Fig. 2b, Fig. S4, Fig. S5).

**Figure 2:**
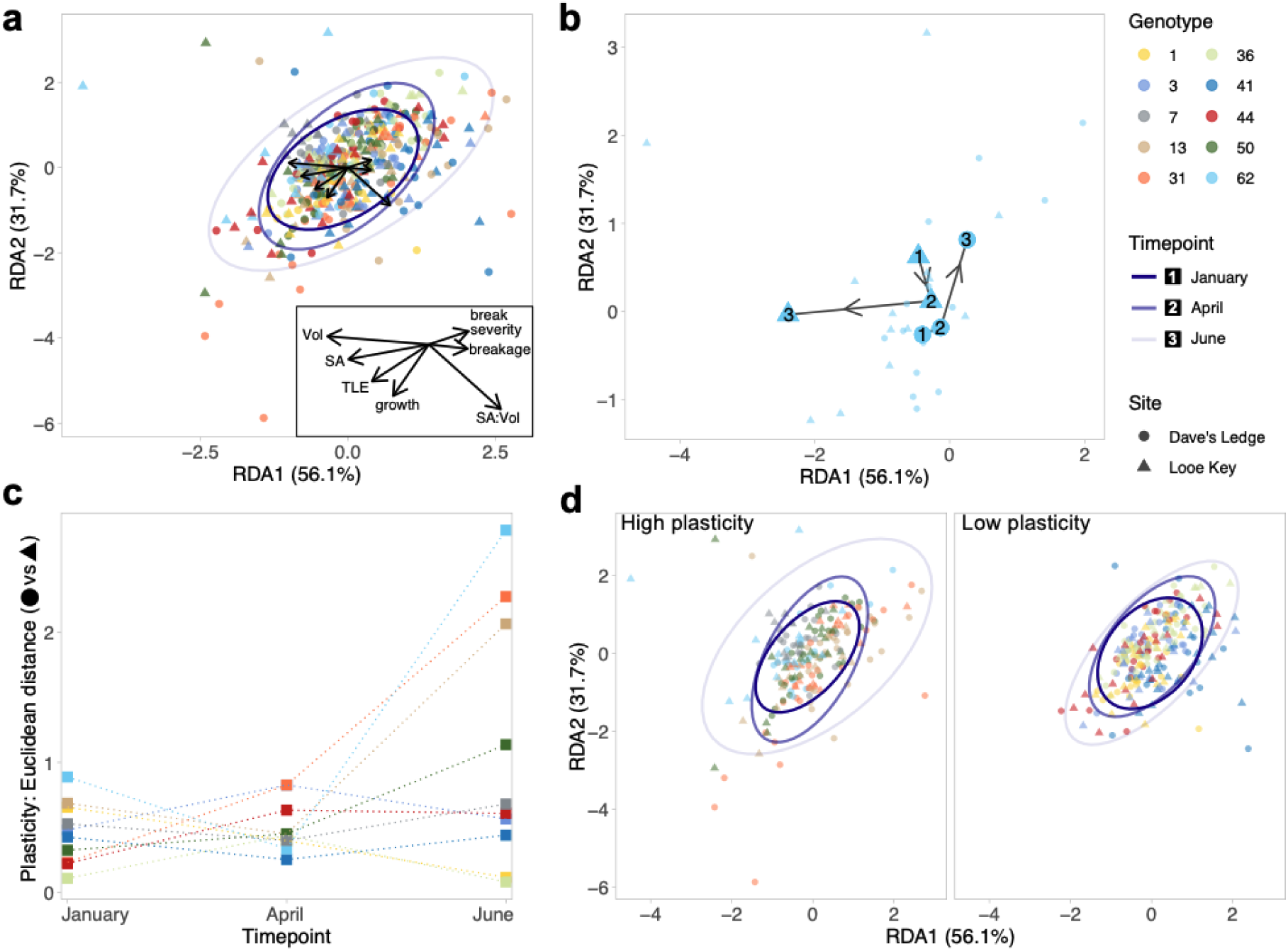
Morphological plasticity is temporally dynamic. Point colors indicate genotype, while shapes represent outplant site. Ellipses show the 95% confidence interval at each timepoint, represented by transparency. RDA plots in panels a, b and d are identical, but vectors along with labeled inset, and timepoint ellipses are omitted in some for visual clarity. **a:** RDA of morphological traits of experimental coral by genotype, site, and their interaction, conditioned on timepoint (January 2023, n = 144; April 2023, n = 141; June 2023, n = 139). Vectors represent morphological traits in RDA space, as labeled in inset. **b:** example of one genotype’s site centroids (enlarged points) movement over time, representing the approach used to calculate the Euclidean distance between sites as a measure of plasticity for each genotype (panel c). Smaller points represent positions of individual ramets. Centroids are labeled by timepoint (1 = January, 2 = April, 3 = June). Segments with arrows represent the movement of centroids in RDA space over time. **c:** Plasticity, calculated as the Euclidean distance between site centroids for each genotype, over time. **d:** RDA of morphological traits, faceted by plasticity at the June timepoint. Left: high plasticity genotypes (top 50th percentile in panel c); right: low plasticity genotypes (bottom 50th percentile in panel c).

To evaluate the effect of genotype and changing plasticity over time on survival under ambient temperature conditions (October 2022-June 2023), Cox proportional hazards models were fit to survival data using the packages coxme (Therneau, 2024) and survival (Therneau et al., 2024).

Pearson’s correlations between color scores measured via ImageJ and S:H obtained via qPCR at each outplant site were used to determine consistency between measures of bleaching stress. To investigate potential tradeoffs between plasticity and responses to thermal stress, linear regressions were used to investigate the relationship between genotype plasticity in June and BSI in July. Significant results were further investigated for statistical robustness in three ways: (1) testing for influential observations using multiple diagnostics (Slope DFBETA, DFFITS, Cook’s distance, Leverage, Studentized residual); (2) sensitivity analysis comparing model outcomes in which individual observations were removed from the model to the full model with all genotypes, and (3) using robust regression analysis in the MASS package (Venables and Ripley, 2002) to investigate the weights assigned to individual observations and compare the slope to our original regression model. All analyses were performed in R (version 4.2.1, R Core Team 2025).

## Results

### Genotype by environment interactions influence morphology

Branching coral morphology is influenced by a balance between growth and breakage (Pratchett et al., 2015), traits which we tracked for nine months in replicate ramets of *A. cervicornis* genets outplanted to natural reef sites in the Lower Florida Keys (Fig. 1a). Logistic regression demonstrated that there were no significant differences in ramets excluded from analyses due to attachment failure by site (p = 0.053), genotype (p = 0.055), or their interaction (p = 1.0). Initial measurements of all morphological traits (TLE, SA, Vol, SA:Vol) indicated baseline differences in size among genotypes prior to outplanting (Table S1), so initial size was accounted for in subsequent analyses.

Linear mixed-effects models showed that linear growth was significantly influenced by initial TLE and also varied by timepoint, genotype, and their interaction, as well as the interaction between genotype and site and the three-way interaction between timepoint, genotype, and site.

Timepoint effects were driven by an accelerating growth rate over time as corals increased in size (Fig. 1e, Table S2). Similarly, timepoint was the dominant effect on absolute size in all measured traits, with all mean trait values increasing over time and varying significantly by timepoint (Fig. 1e, Fig S3, Table S2). Initial size had a significant effect on all traits except volume (Table S2). SA, Vol, and SA:Vol also varied significantly by genotype (Fig. S3, Table S2). To determine the effect of genotype, site, and their interaction on absolute size without the influence of timepoint, we used linear mixed-effects models to investigate their relationship with absolute size at the final timepoint in June. At this timepoint, initial size had a significant effect on all traits except SA:Vol (Table S3). Absolute size in Vol varied significantly by genotype, site, and their interaction, while SA and SA:Vol varied significantly by genotype only (Fig. S3, Table S2). Absolute size in TLE did not vary significantly by any factor (Fig. 1e, Table S3).

Given the diverse effects of genotype, site, and their interaction on individual morphological traits, we used redundancy analysis (RDA) to determine their effect on overall morphology by analyzing an integrated matrix of morphological traits including TLE, SA, Vol, SA:Vol, growth, breakage, and breakage severity while controlling for timepoint and initial TLE. We found that timepoint and initial TLE explained 15.8% of the overall variance in morphology, while genotype, site, and their interaction explained an additional 9.5% of morphological variance. There was a significant effect of genotype (p = 0.001) and genotype x site (p = 0.025) on overall morphology, with the first two RDA axes explaining a significant amount of the constrained variance (RDA1: 56.1% of variance, p = 0.001, RDA2: 31.7% of variance, p = 0.001, Fig. 2a). Ellipses representing the 95% confidence interval at each timepoint show that morphological diversity increased over time, while the directions of trait vectors in RDA space indicate that overall morphology is expressed via a balance between growth and breakage (Fig 2a).

### Genotypic variation in morphological plasticity is temporally dynamic

To further explore the dynamics of morphology over time, we calculated the morphological plasticity of each genotype as the Euclidean distance in RDA space between the two site centroids at each timepoint (Leung et al., 2020) (Fig. 2b, Fig. S4, Fig. S5). We found that plasticity is dynamic over time, with different genotypes having higher or lower relative morphological plasticity at different timepoints (Fig. 2c). Overall, morphological plasticity increased over time, with the highest plasticity values occurring at the final survey timepoint in June and differences in plasticity between genotypes becoming more pronounced (Fig. 2c).

When comparing the top 50% highest plasticity genotypes versus the lowest 50% plasticity genotypes based on the final June timepoint in RDA space, high plasticity genotypes demonstrate greater variation in morphology. This difference is particularly evident along the growth-breakage axis, as indicated by the 95% confidence intervals (Fig. 2a, 2d), indicating that high plasticity genotypes entered more diverse morphospaces than the lower plasticity genotypes. Additionally, higher plasticity genotypes tended to have more within-site variation, or noise, in morphology than the lower plasticity genotypes, particularly at the April and June timepoints as plasticity increased (Fig. S6).

### Plasticity tradeoffs manifest under moderate thermal stress

We investigated whether genotypic differences in morphological plasticity affected survival under non-stressful temperature regimes using Cox proportional hazards models. We used two models to explore the role of (1) genotype and (2) time-dependent plasticity on survival during the first nine months of the outplant experiment, before significant temperature stress was accumulated (Fig. 3a). In the genotype model, only genotype 62 had a significantly higher risk of mortality, by 11.6-fold (p = 0.02, Fig. 3b) compared to the best-surviving reference genotype, 36. The plasticity model showed no significant effect of changing plasticity over time on mortality risk (p = 0.29, Fig 3b).

**Figure 3:**
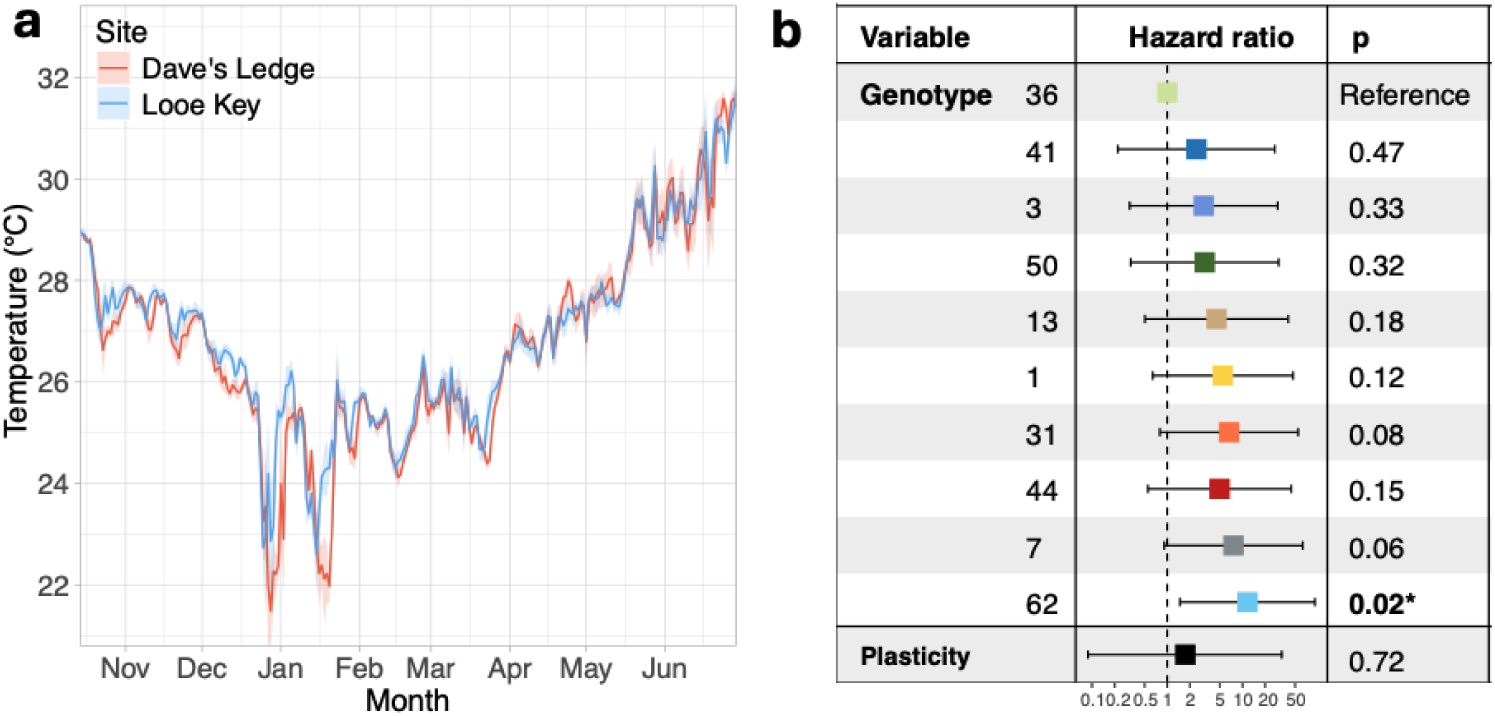
Mortality risk under ambient conditions by genotype and plasticity. **a:** Average daily temperatures in °C (lines) and standard error (shaded area) recorded at the two outplant sites (red = Dave’s Ledge, blue = Looe Key) from outplanting in October 2022 to the final sampling timepoint in June 2023. **b:** Cox proportional hazards ratios (colored squares) for mortality risk by individual genotype (variable “genotype”) with the highest surviving genotype, 36, as the reference; or changing plasticity values over time (variable “plasticity”). Error bars represent 95% confidence intervals. Bolded p-values indicate p<0.05.

At our final regular monitoring timepoint in June, outplants at both sites appeared healthy and did not show visual signs of bleaching stress, although genotype 31 had significantly lighter coloration than genotype 7 (ANOVA, p = 0.00057, Fig. S7), suggesting some ramets may have begun paling. However, beginning mid-June 2023, temperatures began exceeding local bleaching thresholds (“Daily 5km Satellite Regional Virtual Station Time Series Data for Looe Key Reef (v 3.1),” 2025) (Fig. 4a). By late July, we recorded mortality, bleaching, and paling as measured by color scores (Fig. 4b) and S:H ratios (Fig. S8) at both outplant sites. Temperatures were higher at Dave’s Ledge (max = 33.1°C) than at Looe Key (max = 32.3°C) (Fig. 4a), and outplants at Dave’s Ledge accumulated 11.0 degree heating weeks (DHWs) while those at Looe Key accumulated 10.3 DHWs by late July. As a result, there was more widespread mortality, severe bleaching, and greater loss of algal symbionts across genotypes at Dave’s Ledge, and more genotypic variation in bleaching response at Looe Key (Fig. 4b, Fig. S8).

**Figure 4:**
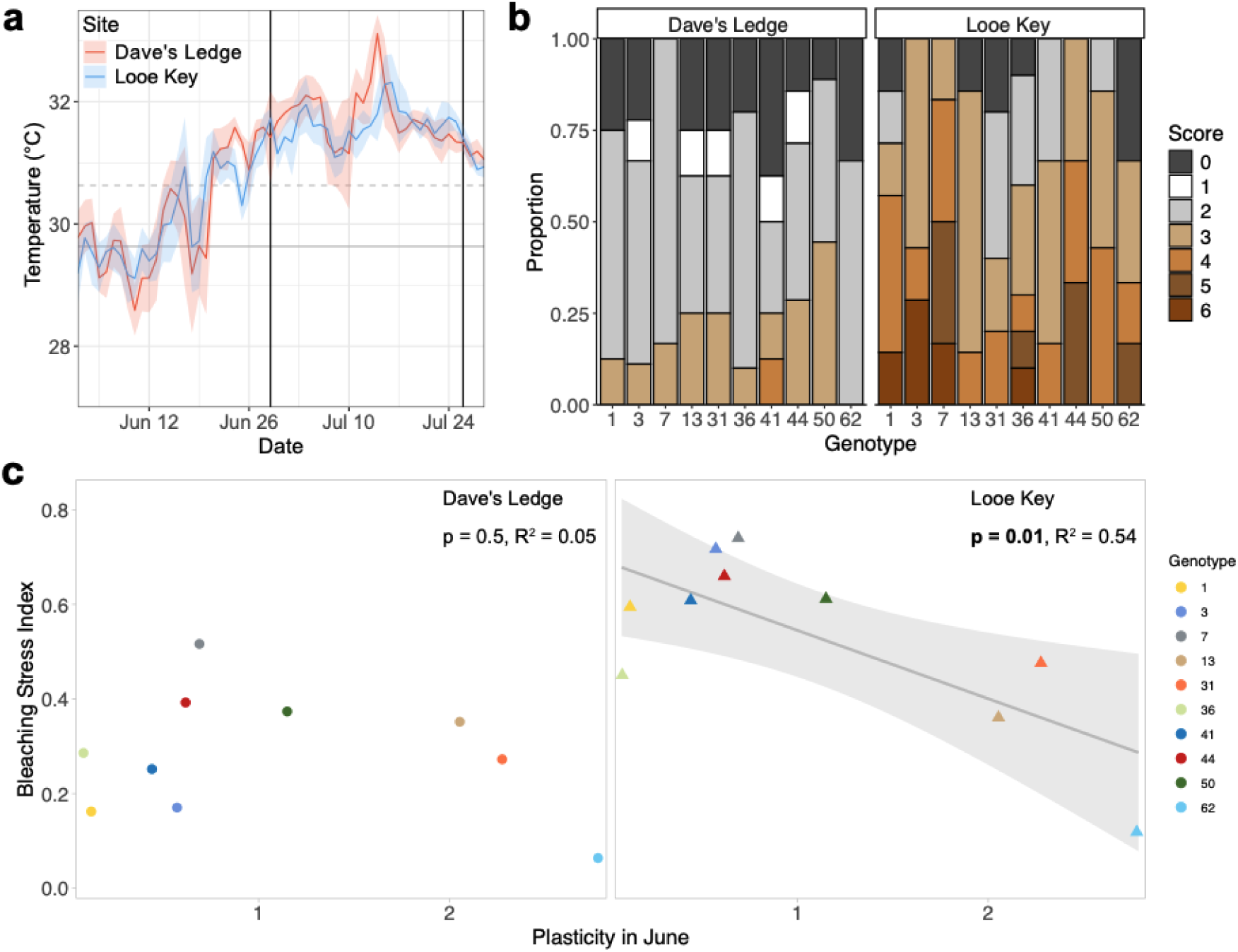
Morphological plasticity trades off with thermal stress resistance. **a:** Average daily temperatures in °C (solid lines) and standard error (shaded areas) measured at the outplant sites immediately prior to and during the bleaching event. Colors represent the outplant site. Black vertical lines represent the June and July monitoring/sampling timepoints. Solid grey horizontal line represents the maximum monthly mean temperature (MMM) for Looe Key Reef (29.63°C)(“Daily 5km Satellite Regional Virtual Station Time Series Data for Looe Key Reef (v 3.1),” 2025), dashed grey horizontal line represents the local bleaching threshold (MMM + 1°C = 30.63°C). **b:** Proportion (y-axis) of surviving experimental corals at the June timepoint (n=139) at each outplant site falling into one of the Coral Health Chart score bleaching (1-6) or mortality (0) categories in July, by genotype along the x-axis. **c:** Relationships between genotype-specific morphological plasticity in June and the Bleaching Stress Index of each genotype at each outplant site in July. For Looe Key, linear regression is represented by the grey line, and the grey shaded area represents the 95% confidence interval. Point colors represent genotype, shapes indicate outplant site.

We integrated genotype bleaching color scores and mortality at the two outplant sites to generate a bleaching stress index (BSI) ranging from 0 (all ramets dead) to 1 (all ramets alive and visually healthy) for each genotype by site combination (Humanes et al., 2024). To investigate whether symbiont community composition may have played a role in bleaching response variation, we used qPCR to check for the presence of thermally tolerant *Durusdinium* (∼*trenchii*) in a subset of samples, but found that across genotypes and sites, most samples were dominated by *Symbiodinium* (∼*fitti*) and only contained background amounts of other symbiont genera (Fig. S8). While some samples did have elevated proportions of *Durusdinium*, they were characterized by very low symbiont concentrations overall (Fig. S8). Additionally, we found no significant correlation (Pearson’s r, p = 0.056) between the proportion of *Durusdinium* detected in an outplant and its color score at the July timepoint (Fig. S8), indicating that symbiont community composition did not play a dominant role in thermal stress response variation.

To investigate potential tradeoffs between genotype-specific morphological plasticity and the response to thermal stress, we explored the relationships between plasticity values measured at the June monitoring timepoint and genotype BSIs at the two outplant sites in July. At Dave’s Ledge, where temperature stress was more extreme and genotypic variation in bleaching response was lower (Fig. 4a, 4b; Fig. S8), there was no relationship between plasticity in June and the response to thermal stress (p = 0.38, Fig 4c). However, at Looe Key, where temperatures were slightly lower and more genotypic variation in the bleaching response remained (Fig 4a, 4b), there was a significant negative relationship between morphological plasticity and BSI (p = 0.010, Fig 4c) with more plastic genotypes showing more negative bleaching outcomes. Although two observations at Looe Key (genotypes 36 and 62) were flagged as influential in our analyses (Table S5), a sensitivity analysis where individual genotypes were omitted from the model showed a consistent negative relationship between plasticity and bleaching outcomes regardless of which observation was excluded from the model, although the model without genotype 62 was no longer statistically significant (Table S6). Furthermore, a robust regression analysis resulted in a very similar slope estimate as our original ordinary regression model (ordinary: -0.144; robust: -0.147), with all genotypes being weighted >0.9 with the exception of genotype 36 (Table S7). Together, these results indicate that despite two influential observations, the data support a consistent and statistically significant negative relationship between genotypic plasticity and bleaching outcomes at Looe Key.

## Discussion

Phenotypic plasticity may play an outsize role in the persistence of sessile, long-lived organisms under the rapid environmental shifts occurring due to climate change due to their unique life-history (Chevin et al., 2010; Kelly, 2019). However, plasticity is often associated with costs and limits (DeWitt et al., 1998) which can be hard to detect (Auld et al., 2009). Here, we leveraged clonal reproduction via fragmentation in *A. cervicornis* to reveal reduced thermal stress resistance at more moderate temperature stress levels as one consequence of elevated morphological plasticity in this critically endangered coral species. Though the unprecedented 2023 marine heatwave captured during this experiment (Hoegh-Guldberg et al., 2023; Spady et al., 2026) meant we could no longer detect genotypic variation in thermal stress resistance at one of our sites, we found that increased resistance to bleaching and mortality was negatively correlated with morphological plasticity at the site which experienced lower levels of temperature stress. While previous work in this system reported no tradeoffs between plasticity and survival or growth under ambient conditions (Million et al., 2022), we found that morphological plasticity was neutral under non-stressful conditions preceding the bleaching event and maladaptive under moderate thermal stress, illustrating context-dependent consequences of plasticity. Additionally, we observed a negative tradeoff where the highest plasticity genotypes were the most sensitive to thermal stress at Looe Key, aligning with previous work which found that costs of plasticity tend to be strongest when plastic responses are large (Hendry, 2016). For example, a study in the common frog (*Rana temporaria*) only found costs of plasticity in development time under dry conditions in populations with the largest plastic responses (Lind and Johansson, 2009). Our results suggest that *A. cervicornis* could not maximize both plasticity and thermal tolerance in our experiment, meaning the benefits of morphological plasticity could be constrained by costs and/or limits (DeWitt et al., 1998).

While this field-based transplant experiment was limited to 10 genotypes outplanted at two reef sites, Florida’s *A. cervicornis* population is genetically well-mixed (Duffin et al., 2026). Moreover, previous work has shown that sampling 10 genets is sufficient to capture common allelic diversity of *A. cervicornis* in Florida (Drury et al., 2017) and the genotypes selected for this study are broadly representative of genetic diversity in the population (Fig. S1). Additionally, the two outplant sites selected for this experiment were informed by the intermediate survivorship at these sites in a previous study which outplanted these same 10 genotypes across 9 sites in the Lower Florida Keys (Million et al., 2022), suggesting they are representative of non-stressful environmental conditions for these outplants. It is also important to note that this population has been severely reduced by the impacts of climate change, leading to the functional extinction of *A. cervicornis* from Florida’s coral reefs (Manzello et al., 2025). As a result, the genotypes that are maintained in captivity by restoration practitioners are instrumental for preventing the extirpation of these species (Muller et al., 2025), and will play a key role in conservation and restoration going forward (Suggett et al., 2024). Thus, this representative study of 10 genotypes that are widely used in Florida’s restoration efforts offers insights into the broader eco-evolutionary dynamics of interactions between morphological plasticity and thermal stress tolerance in *A. cervicornis*.

In this study, we find that morphological plasticity in *A. cervicornis* is expressed as a balance between growth and breakage, so one possible cost of plasticity is the high energetic investment required for growth (Cohen and McConnaughey, 2003). While tradeoffs between coral thermal tolerance and growth have been demonstrated in several species, they are usually associated with major differences in the community composition of endosymbiotic algae (Little et al., 2004). Under ambient conditions, coral hosting symbionts in the more thermally tolerant genus *Durusdinium* may have reduced growth rates compared to conspecifics hosting the more thermally sensitive *Cladocopium* (Cunning et al., 2015), as *Durusdinium* endosymbionts translocate less carbon to the host, resulting in less energy available for growth (Cantin et al., 2009). In species with less variation in their algal symbiont communities, such as *A. cervicornis* in our experiment, evidence for tradeoffs between growth and thermal tolerance is mixed.

Positive associations between surface area and volumetric growth and thermal tolerance were observed in an Indo-Pacific congener, *Acropora digitifera* (Lachs et al., 2023), while a prior study in *A. cervicornis* found a genotype-mediated negative association between linear growth rates before thermal stress and percent tissue loss after bleaching (Ladd et al., 2017). The lack of a consistent relationship between growth and thermal tolerance in species with more uniform endosymbiont communities indicates that tradeoffs may be more subtle or involve growth-related plasticity. Particularly, associations between individual growth metrics and fitness traits may not be sufficient to detect potential tradeoffs. Capturing and integrating multiple growth metrics to track morphological plasticity over time allowed us to reveal the context-dependent tradeoff of plasticity with thermal tolerance at moderate temperatures in this study, demonstrating the need to incorporate more holistic considerations of growth when measuring tradeoffs with thermal tolerance.

The observation that high plasticity genotypes express more varied morphologies via a balance of growth and breakage suggests that the mechanistic tradeoff with thermal tolerance could occur via production costs, or the cost to implement a plastic response (DeWitt et al., 1998). In *A. cervicornis*, morphological plasticity is modulated in part by calcification-mediated skeletal extension (Pratchett et al., 2015) and potentially the complementary process of skeletal infilling (Gladfelter, 1982), which influences structural integrity and propensity to fragment (Highsmith, 1982; Tunnicliffe, 1981). Although the precise mechanism remains unresolved, multiple studies have produced evidence in support of active ion transport being involved in the formation of new skeleton (Allemand et al., 2011; Cohen and McConnaughey, 2003). Moreover, bleaching, the visual manifestation of starvation via the loss of photosynthate from algal endosymbionts, is known to yield long-term calcification deficits (Rodrigues and Grottoli, 2006). Baseline differences among genotypes in skeletal extension or infill rates could result in increased investment in these energy-intensive processes or the need to recover from more frequent breakage, resulting in less energy stores available for responses to thermal stress.

In addition to revealing context-dependent costs, we find that morphological plasticity in *A. cervicornis* is dynamic over time: different genotypes had higher relative plasticity at different timepoints throughout our study. Since the balance between growth and breakage is vital to the expression of morphological plasticity, we hypothesize that the relative plasticity of different genotypes changes over time as corals increase in size and complexity, yielding more branches along which growth and breakage can occur. Additionally, the first nine months of outplant morphology tracked during this experiment represent a substantial change in morphological complexity, as simple ramets with no branches transformed into more complex colonies. It is possible that this study characterizes a highly dynamic phase in plasticity at the early stages of coral growout, and that the relative plasticity of the genotypes studied here could stabilize over time once ramets reach a certain size and their complexity is no longer as strongly in flux.

In addition to temporal variation in our focal morphological traits, within-site morphological variation, or noise, between ramets of the highest plasticity genotypes was also elevated, particularly at the later timepoints when morphological complexity was greatest. This positive association between within-site noise and between-site plasticity is indicative of a trend that has previously only been demonstrated in model systems. In yeast, the relationship between noise and plasticity in gene expression is strongly coupled (Lehner, 2010), and in *Drosophila*, developmental noise and phenotypic plasticity in wing morphology are positively correlated (Saito et al., 2024). While the underlying mechanisms leading to the genetic correlation between noise and plasticity remain poorly understood, our observations suggest they are also ecologically important.

The dynamic fluctuation of morphological plasticity in these coral outplants over time indicates that phenotypic changes are likely influenced by many environmental variables, meaning that the adaptive value of morphological plasticity in one environment or at one particular timepoint may differ from other environmental or temporal contexts. This is supported by the contrasting results between this study, which finds a neutral effect of time-dependent plasticity at non-stressful temperatures, and a previous study of the same genotypes across a larger suite of outplant sites, which found that morphological plasticity was adaptive at non-bleaching temperatures (Million et al., 2022). Similar patterns of environmentally dynamic plasticity were found in montane butterflies, where plasticity in wing absorptivity, which is important for adaptive thermal regulation, varied by elevation (Kingsolver and Buckley, 2017). As such, while we can consider the adaptive value of plasticity in certain contexts, whether it is adaptive overall requires assessment in multiple environments (Hendry, 2016). Additionally, it is important to consider temporal dynamics in the context of life-history. In many other systems, temporal dynamics may only become important in a trans-generational context (Fox et al., 2019), but in long-lived clonal organisms it is essential to consider the temporal dynamics that may occur within one individual’s lifetime.

Caribbean acroporids, including *A. cervicornis,* are highly fragmentation-prone species, where asexual clonal ramets regularly establish populations in new habitats (Drury et al., 2019; Highsmith, 1982). Similarly, ramets of the same genotypes are placed in many different reef environments for the purposes of reef restoration (Young et al., 2012). The capacity for morphological plasticity may therefore have been historically advantageous, as plasticity may be beneficial in establishing populations in new habitats. For example, a meta-analysis of 75 invasive and noninvasive plant species pairs found that invading species had greater plasticity (Davidson et al., 2011). However, even if morphological plasticity evolved in *A. cervicornis* due to its neutral to adaptive fitness effects under historical environmental conditions, more plastic genotypes could now be at a disadvantage more often under contemporary climate-driven conditions as increasing temperatures and more frequent and severe marine heatwaves (Hughes et al., 2017) expose them to environmental conditions under which these context-dependent tradeoffs are revealed.

The implications of contemporary marine heatwaves are further underscored by the fact that although there was evidence of genotypic variation in the response to thermal stress at Looe Key early in the 2023 mass bleaching event, corals at Dave’s Ledge were exposed to much more severe thermal stress, resulting in widespread bleaching and mortality regardless of genotype. Additionally, the unprecedented magnitude and duration of the 2023 marine heatwave in the Florida Keys meant that the genotypic variation in bleaching response we observed at one of our sites in July was overwhelmed by extreme thermal stress of over 20 DHWs (Hoegh-Guldberg et al., 2023; Spady et al., 2026), and almost all of the outplants in this experiment were dead by November 2023 (Fig. S9). Such high levels of mortality were widespread throughout the region during the 2023 marine heatwave, resulting in the functional extinction of *A. cervicornis* in Florida (Manzello et al., 2025). While our results suggest that the context-dependent tradeoff between moderate thermal stress resistance and plasticity could constrain the evolution of morphological plasticity in *A. cervicornis* and may have played a role in the maintenance of genotypic variation in plasticity under historic temperature regimes, contemporary thermal stress events have already swamped the adaptive capacity of this key reef-building species.

## Acknowledgments

We are grateful to Erich Bartels, Zachary Craig, Kyle Knoblock, Amanda Lewan, Samantha Simpson, and Cory Walter at Mote Marine Laboratory and Sibelle O’Donnell at USC for their assistance with fieldwork. Coral were outplanted and sampled under FKNMS permit 2022-120. Funding for this study was provided by NSF IOS 2222272 to CDK and NSF IOS 2222273 to HRK/EMM.

## Author contributions

JD: contributed to study design, performed field research, developed methodology, processed molecular samples, and collected, curated, analyzed, and visualized data. MG: performed field research, developed methodology, and collected and curated data. IC: contributed to funding acquisition and study design and performed field research. IH: collected and curated morphological data. JK: performed and coordinated field research. SL: performed field research and collected and curated morphological data. TV: collected and curated morphological data. NV: processed molecular samples. HRK: contributed to funding acquisition and study design and provided resources. EMM: contributed to funding acquisition and study design and provided resources. CDK: designed the study, acquired funding, and provided supervision and resources. JD wrote the first draft of the manuscript, and all authors contributed to manuscript review and editing.

## Statement on inclusion

Our study arises from a collaborative research initiative, bringing together authors based at the University of Southern California in Los Angeles, California and Mote Marine Laboratory’s Elizabeth Moore Center in Summerland Key, Florida, where the fieldwork component of this study was carried out.

## Data availability statement

All data and analytical code are available at https://doi.org/10.5281/zenodo.20346481. Workflow and scripts for batch processing 3D models can be found at https://doi.org/10.5281/zenodo.17136226.

