## Supplemental Information for "Context-dependent tradeoffs between morphological plasticity and thermal stress resistance in a reef-building coral"

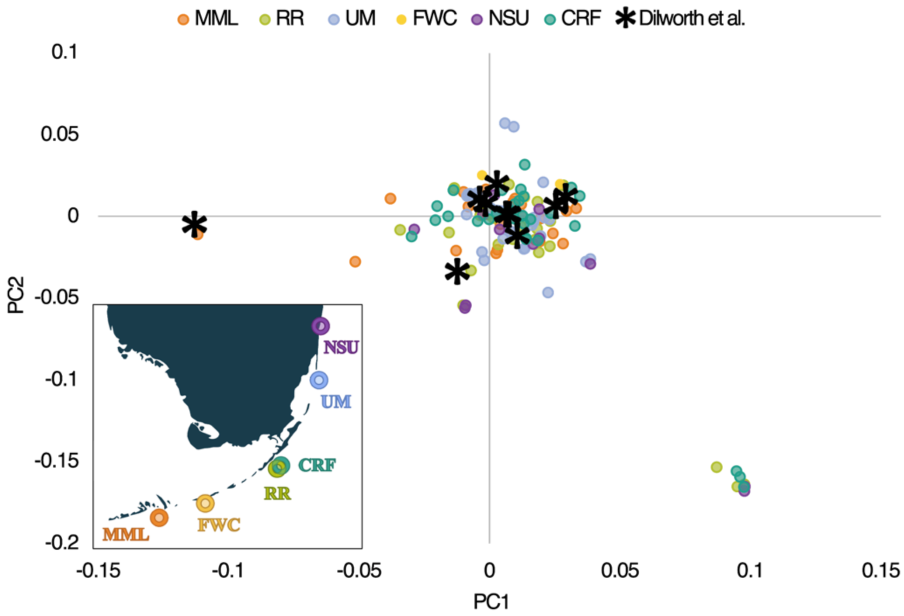


**Figure S1:** PCA showing genomic variation revealed through whole genome sequencing of 133 unique *A. cervicornis* genotypes sampled from restoration nurseries throughout the Florida Reef Tract (data from Duffin et al., 2026). Point colors indicate source (MML = Mote Marine Lab, RRT = Reef Renewal, UM = University of Miami, FWC = Florida Fish and Wildlife Conservation Commission, NSU = Nova Southeastern University, CRF = Coral Restoration Foundation). Genotypes used in this study are represented by stars. Inset map shows geographic locations of sampled nurseries along the Florida Reef Tract.

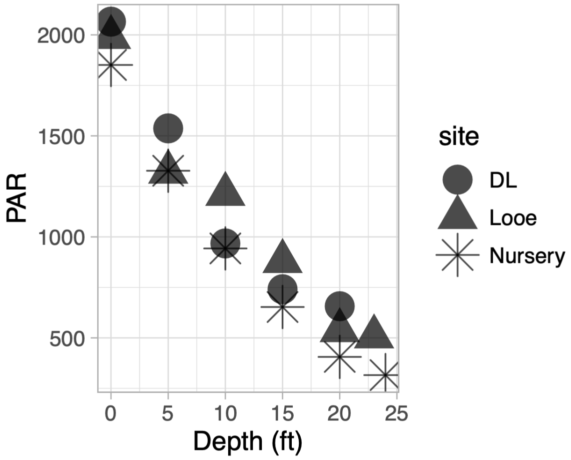


**Figure S2:** Photosynthetic active radiation (PAR) measured across depths at the two outplant sites (DL = Dave's Ledge, Looe = Looe Key) and the source nursery on April 18th and 20th 2023. Measurements were taken within 30 minutes of high noon, and weather conditions were cloudless on both measurement days. Shapes represent measurement site.

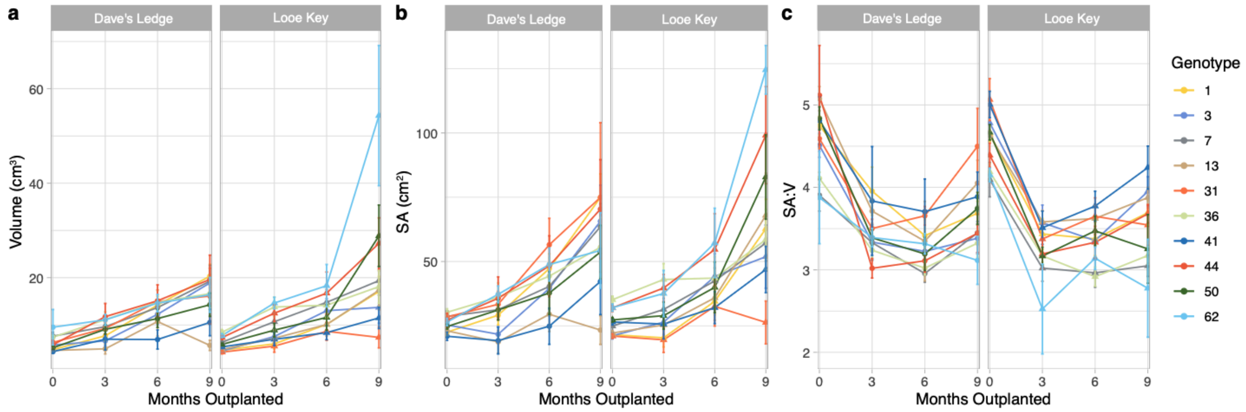


**Figure S3:** Genotype mean volume in cm$^3$ (a), surface area in cm$^2$ (b) and surface area:volume ratio (c) at each outplant site from initial (October 2022, n = 159) through the following three-month monitoring intervals (January 2023, n = 144; April 2023, n = 141; June 2023, n = 139). Point colors represent genotype, shapes represent site. Error bars show standard error.

**Table S1.** Results from two-way ANOVAs testing for baseline differences in morphological traits by genotype and outplant site. Bold font indicates significant p-values.

| **Trait** | **Factor** | **F value** | **p-value** |
|---|---|---|---|
| TLE | genotype | 9.247 | **6.47x10$^{-11}$** |
| | site | 0.812 | 0.369 |
| | genotype*site | 4.678 | **1.37x10$^{-8}$** |
| SA | genotype | 11.206 | **4.74x10$^{-13}$** |
| | site | 0.874 | 0.351 |
| | genotype*site | 4.755 | **1.59x10$^{-5}$** |
| Vol | genotype | 6.179 | **2.65x10$^{-7}$** |
| | site | 0.130 | 0.719 |
| | genotype*site | 0.1296 | 0.244 |
| SA:Vol | genotype | 5.106 | **5.74x10$^{-6}$** |
| | site | 0.007 | 0.931 |
| | genotype*site | 1.277 | 0.255 |

**Table S2**: Type III ANOVA results (Satterthwaite's method) of linear mixed-effects models investigating the effect of initial size, as well as timepoint, genotype, site, and their interactions on growth and size metrics. Tag number was included as random intercept in all models. Asterisks and bolded text indicate significance levels.

| **Metric** | **Factor** | **F-value** | **p-value** |
|---|---|---|---|
| growth (positive change in TLE) | Initial TLE | 7.2485 | **0.008284 **** |
| | timepoint | 74.4754 | **<2.2 x10$^{-16}$ ***** |
| | genotype | 3.3453 | **0.001288 **** |
| | site | 0.1584 | 0.691522 |
| | timepoint*genotype | 2.5694 | **0.001009 **** |
| | timepoint*site | 0.5006 | 0.607167 |
| | genotype*site | 3.0281 | **0.003093 **** |
| | timepoint*genotype*site | 3.5752 | **8.661x10$^{-6}$ ***** |
| TLE | Initial TLE | 7.1020 | **0.008859 **** |
| | timepoint | 20.0737 | **1.103-x10$^{-8}$ ***** |
| | genotype | 1.8290 | 0.070881 |
| | site | 1.3493 | 0.247938 |
| | timepoint*genotype | 1.3821 | 0.143089 |
| | timepoint*site | 1.5245 | 0.220208 |
| | genotype*site | 0.4923 | 0.876976 |
| | timepoint*genotype*site | 0.6621 | 0.845582 |
| Vol | Initial Vol | 3.5304 | 0.064599 |
| | timepoint | 6.3590 | **0.002223 **** |
| | genotype | 2.1239 | **0.038241 *** |
| | site | 1.2460 | 0.268037 |
| | timepoint*genotype | 0.8080 | 0.688767 |
| | timepoint*site | 1.5176 | 0.222509 |
| | genotype*site | 0.7031 | 0.703978 |
| | timepoint*genotype*site | 1.1601 | 0.301317 |

| | | | |
|---|---|---|---|
| SA | Initial SA | 19.5683 | **$2.269 \times 10^{-5}$ ***** |
| | timepoint | 20.2430 | **$9.543 \times 10^{-9}$ ***** |
| | genotype | 2.5591 | **0.01036 *** |
| | site | 0.0094 | 0.92307 |
| | timepoint*genotype | 0.8235 | 0.67138 |
| | timepoint*site | 2.4210 | 0.09138 |
| | genotype*site | 0.7200 | 0.68964 |
| | timepoint*genotype*site | 1.3388 | 0.16664 |
| SA:Vol | Initial SA:Vol | 4.3489 | **0.0392738 *** |
| | timepoint | 9.3448 | **0.0001263 ***** |
| | genotype | 6.2734 | **$3.157 \times 10^{-7}$ ***** |
| | site | 1.2135 | 0.1634441 |
| | timepoint*genotype | 1.1853 | 0.2513756 |
| | timepoint*site | 2.3957 | 0.0934257 |
| | genotype*site | 0.8452 | 0.5761708 |
| | timepoint*genotype*site | 0.8373 | 0.6550816 |

**Table S3:** Type III ANOVA results (Satterthwaite's method) of linear mixed-effects models investigating the effect of initial size, as well as genotype, site, and their interaction on size metrics at the final measurement timepoint in June. Outplant array was included as a random intercept in all models. Asterisks and bolded text indicate significance levels.

| Metric | Factor | F-value | P-value |
|---|---|---|---|
| TLE | Initial TLE | 12.9252 | **0.0004845 ***** |
| | genotype | 1.5082 | 0.1546567 |
| | site | 1.6976 | 0.2122432 |
| | genotype*site | 1.5818 | 0.1303205 |
| Vol | Initial Vol | 8.8203 | **0.003645 **** |
| | genotype | 3.2494 | **0.001525 **** |
| | site | 10.5408 | **0.001541 **** |
| | genotype*site | 4.3446 | **$7.145 \times 10^{-5}$ ***** |
| SA | Initial SA | 22.9296 | **$5.182 \times 10^{-6}$ ***** |
| | genotype | 2.2833 | **0.02167 *** |
| | site | 1.8719 | 0.17400 |
| | genotype*site | 1.4687 | 0.16819 |
| SA:Vol | Iniital SA:Vol | 0.7719 | 0.381520 |
| | genotype | 3.3564 | **0.001131 **** |
| | site | 1.3484 | 0.248035 |
| | genotype*site | 1.4379 | 0.180430 |

**Table S4:** Combinations of breakage type and number and net growth resulting in final breakage severity categories used in RDA matrix. P = primary branch, S = secondary branch, T = tertiary branch. Catastrophic breakage indicates loss of all higher-level branches and most of the primary branch.

| Breakage Type/Number | Net Growth | Breakage Severity |
|---|---|---|
| none | Growth > 0 | 0 |
| P, S, T breaks adding up to 1 | Growth > 0 | 1 |
| P, S, T breaks adding up to 2 | N/A | 2 |
| P, S, T breaks adding up to 3 | N/A | 3 |
| P, S, T breaks adding up to 4 | Growth < 0 | 4 |
| Catastrophic breakage | Growth < 0 | 5 |

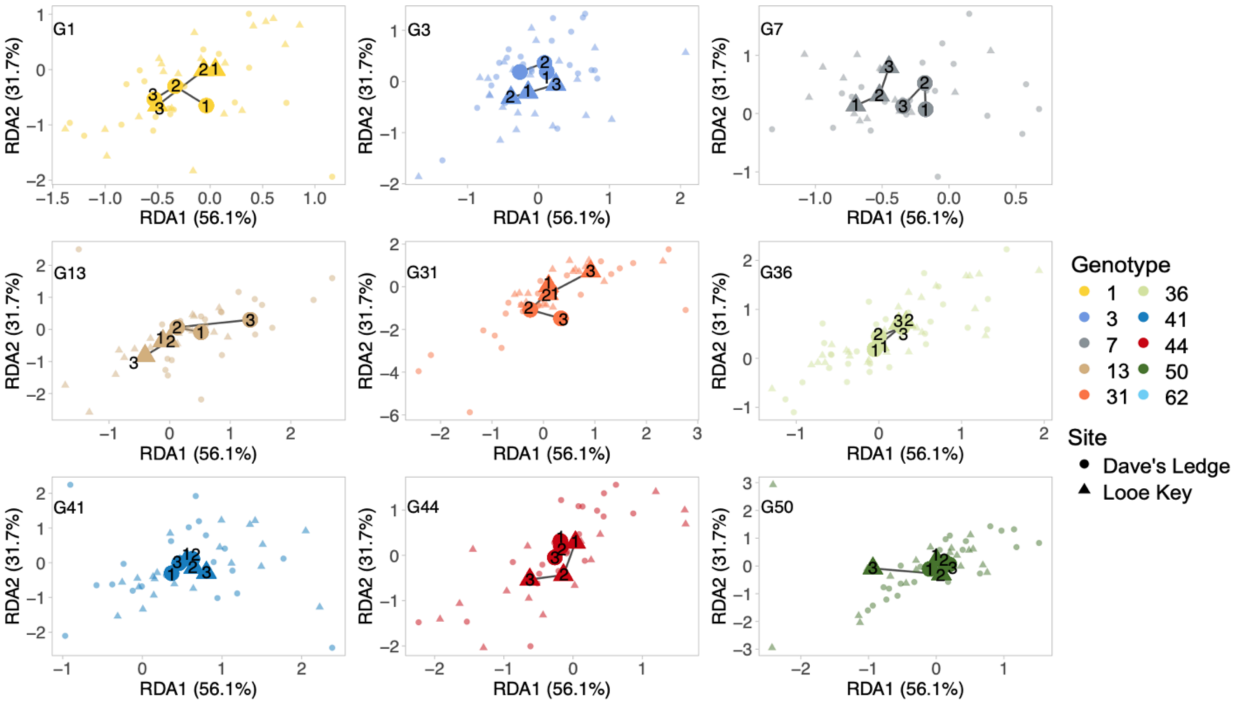


**Figure S4:** RDA of morphological traits of experimental coral by genotype, site, and their interaction, conditioned on timepoint, faceted by genotype. Enlarged points show each genotype's site-specific centroid movement over time, representing the approach used to calculate the Euclidean distance between sites as a measure of plasticity for each genotype (Fig. 2c). Colors represent genotype, shapes represent outplant site.

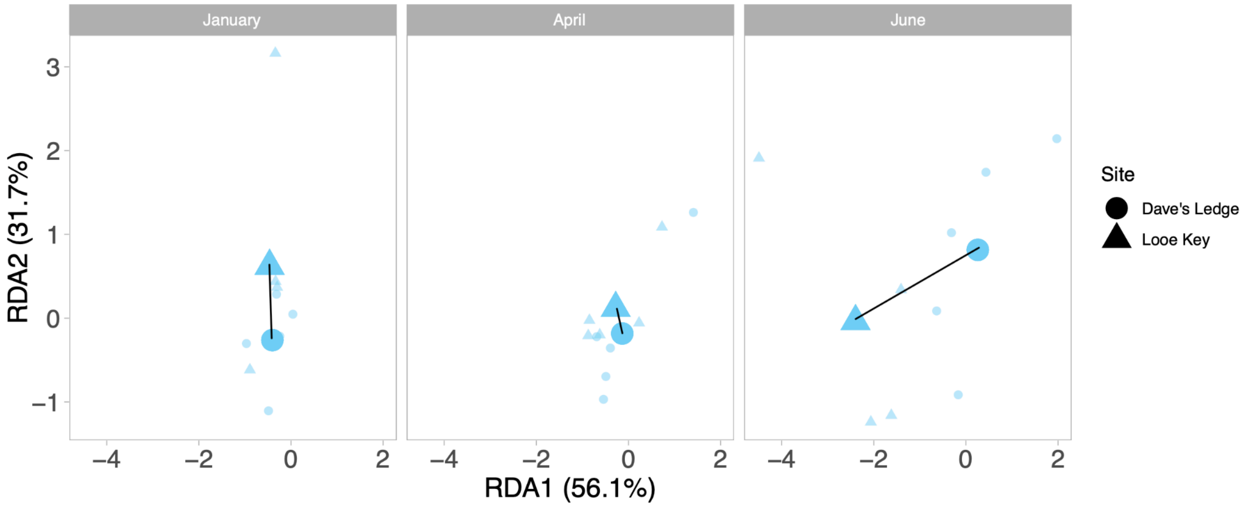


**Figure S5:** Illustrative example of genotype 62, representing the approach used to calculate plasticity (Fig 2c). Plasticity was measured as the Euclidean distance between site-specific centroids (enlarged points) at each faceted timepoint. Smaller points represent positions of individual ramets. Shapes represent outplant site, and black lines represent the Euclidean distance measured between site centroids at each timepoint.

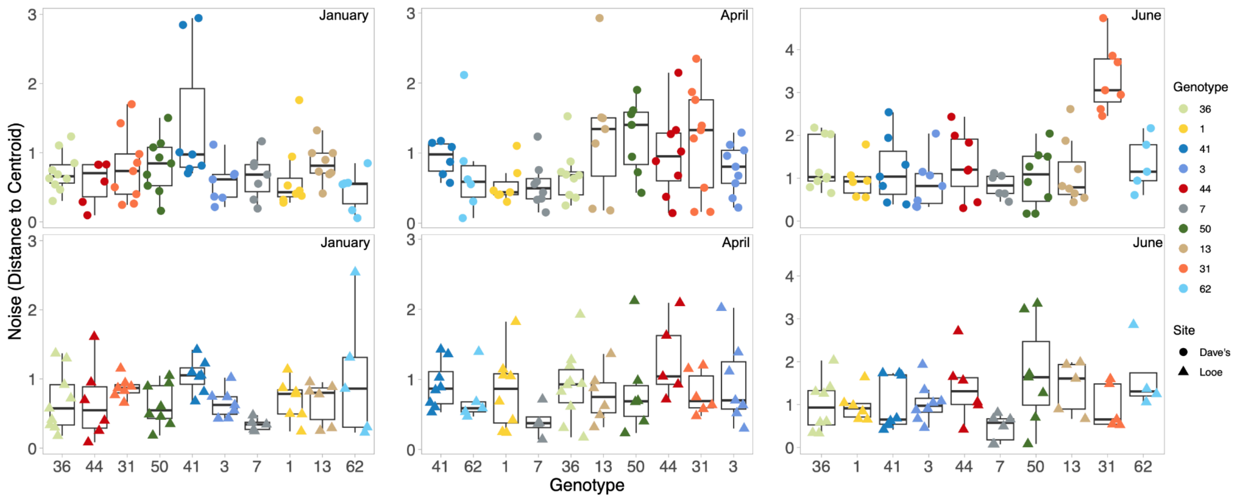


**Figure S6:** Boxplots showing the distribution of noise, calculated as the Euclidean distance between each sample and the site centroid for each genotype at each measurement timepoint. Genotypes on the x-axis are ordered from highest to lowest plasticity at each timepoint. Top row: Dave's Ledge, bottom row: Looe Key. Point colors represent genotype, shapes represent site.

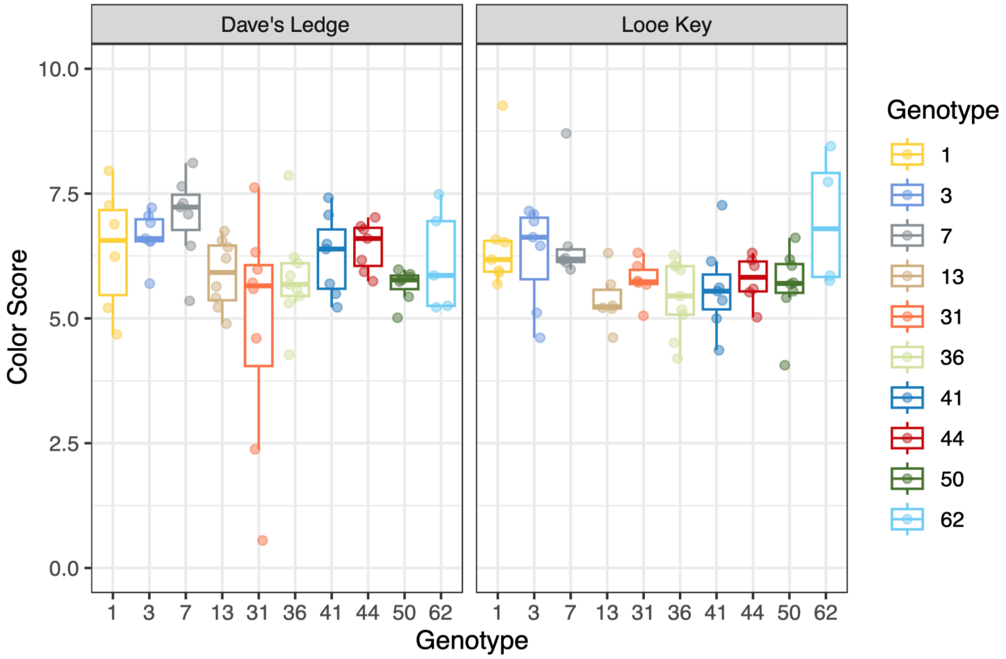


**Figure S7:** Boxplot of Coral Health Chart color scores measured at the final June sampling timepoint before the onset of severe thermal stress, faceted by outplant site, genotype along the x-axis and represented by color.

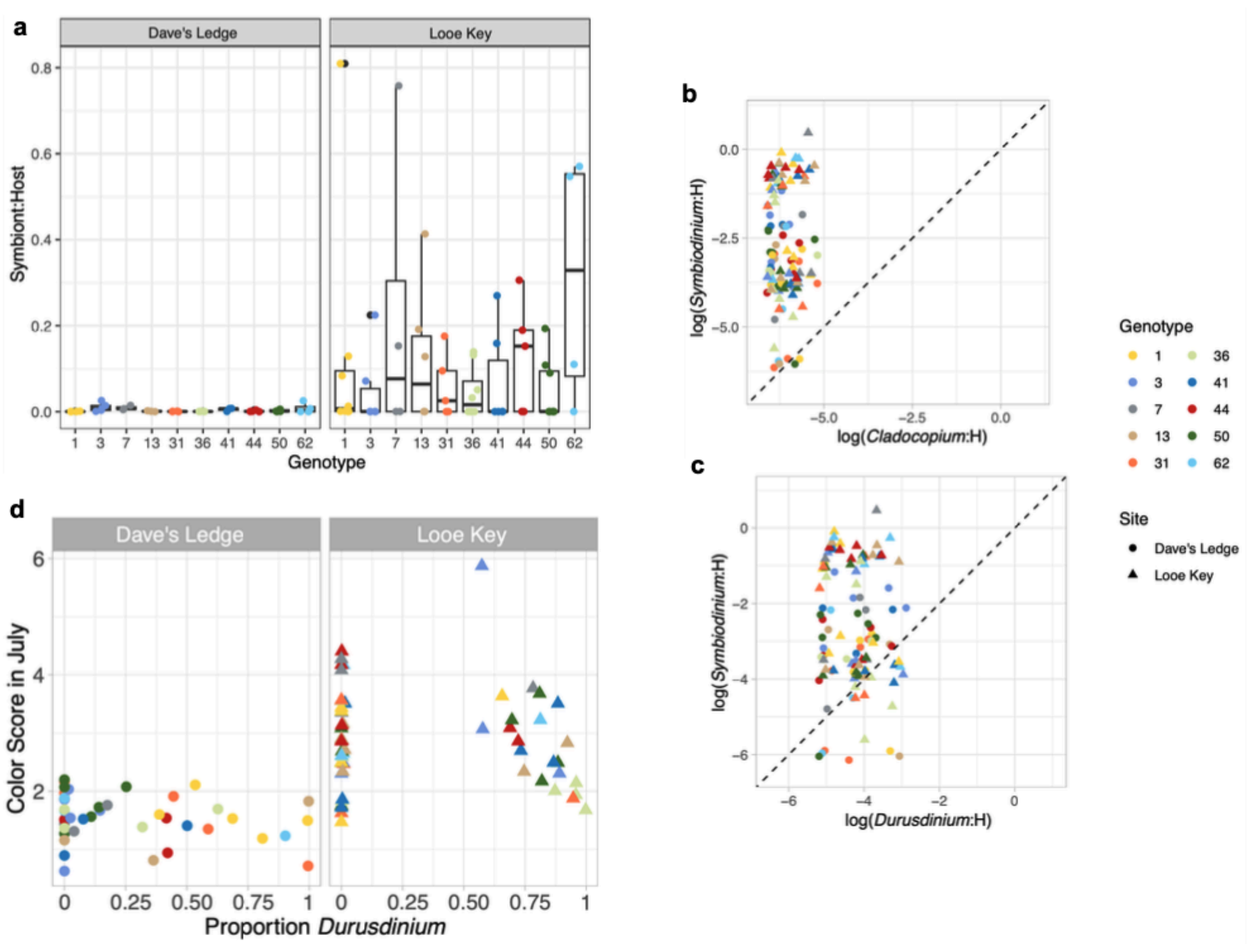


**Figure S8: a:** Boxplot of symbiont:host ratios measured via qPCR from total DNA extracted from samples collected at the July sampling timepoint during the bleaching event. Faceted by outplant site, genotype along the x-axis and represented by color. **b-c:** Base 10 log of *Symbiodinium*-to-host ratio against base 10 log of *Cladocopium*-to-host ratio (b) and *Durusdinium*-to-host ratio (c). Shapes represent outplant site, colors indicate genotype. Dashed diagonal lines represent a 1:1 ratio of *Symbiodinium* to *Cladocopium* or *Durusdinium* symbionts. **d:** Scatterplots displaying the relationship between the proportion of thermally tolerant *Durusdinium* symbionts detected in each outplant and its color score in July. Shapes represent outplant site, colors indicate genotype.

**Table S5**: Results of influential observation diagnostics for each of the genotypes included in the linear regression model for Looe Key.

| genotype | Slope DFBETA | DFFITS | Cook's distance | Leverage | Studentized residual | Influential |
|---|---|---|---|---|---|---|
| 36 | 0.997 | -1.352 | 0.541 | 0.219 | -2.550 | true |
| 41 | 0.039 | -0.068 | 0.003 | 0.149 | -0.163 | false |
| 3 | -0.149 | 0.304 | 0.049 | 0.131 | 0.782 | false |
| 50 | 0.013 | 0.198 | 0.021 | 0.100 | 0.593 | false |
| 13 | -0.091 | -0.124 | 0.009 | 0.218 | -0.234 | false |
| 44 | -0.085 | 0.187 | 0.019 | 0.126 | 0.492 | false |
| 1 | 0.205 | -0.283 | 0.044 | 0.211 | -0.548 | false |
| 31 | 0.474 | 0.595 | 0.178 | 0.273 | 0.969 | false |
| 7 | -0.192 | 0.482 | 0.106 | 0.119 | 1.312 | false |
| 62 | -1.489 | -1.687 | 1.090 | 0.452 | -1.855 | true |

*Note: Influential observations were identified using influence.measures() and commonly used diagnostic screening thresholds.*

**Table S6**: Model parameters of individual "leave one out" models from the sensitivity analysis compared to the full Looe Key linear regression model including all genotypes.

| Model | Estimate | Standard error | 95% CI | P value | $R^2$ |
|---|---|---|---|---|---|
| **Full model** | **-0.144** | **0.047** | **-0.252 to -0.037** | **0.015** | **0.545** |
| Without genotype 36 | -0.180 | 0.039 | -0.271 to -0.089 | 0.002 | 0.757 |
| Without genotype 41 | -0.146 | 0.051 | -0.267 to -0.025 | 0.024 | 0.538 |
| Without genotype 3 | -0.137 | 0.049 | -0.252 to -0.022 | 0.026 | 0.532 |
| Without genotype 50 | -0.145 | 0.049 | -0.260 to -0.030 | 0.021 | 0.559 |
| Without genotype 13 | -0.140 | 0.053 | -0.266 to -0.014 | 0.034 | 0.496 |
| Without genotype 44 | -0.140 | 0.050 | -0.258 to -0.022 | 0.026 | 0.531 |
| Without genotype 1 | -0.154 | 0.052 | -0.278 to -0.031 | 0.021 | 0.556 |
| Without genotype 31 | -0.167 | 0.052 | -0.290 to -0.043 | 0.015 | 0.593 |
| Without genotype 7 | -0.136 | 0.045 | -0.243 to -0.029 | 0.020 | 0.563 |
| Without genotype 62 | -0.084 | 0.052 | -0.207 to 0.040 | 0.155 | 0.267 |

*Note: Estimate is the regression slope. Each sensitivity model excludes one genotype. The shaded row represents the model containing all genotypes.*

**Table S7:** Weights assigned to each genotype observation in the Looe Key linear regression model using robust linear regression from the MASS package.

| genotype | Robust weight |
|---|---|
| 36 | 0.792 |
| 41 | 0.997 |
| 3 | 0.966 |
| 50 | 0.979 |
| 13 | 0.996 |
| 44 | 0.987 |
| 1 | 0.978 |
| 31 | 0.954 |
| 7 | 0.913 |
| 62 | 0.904 |

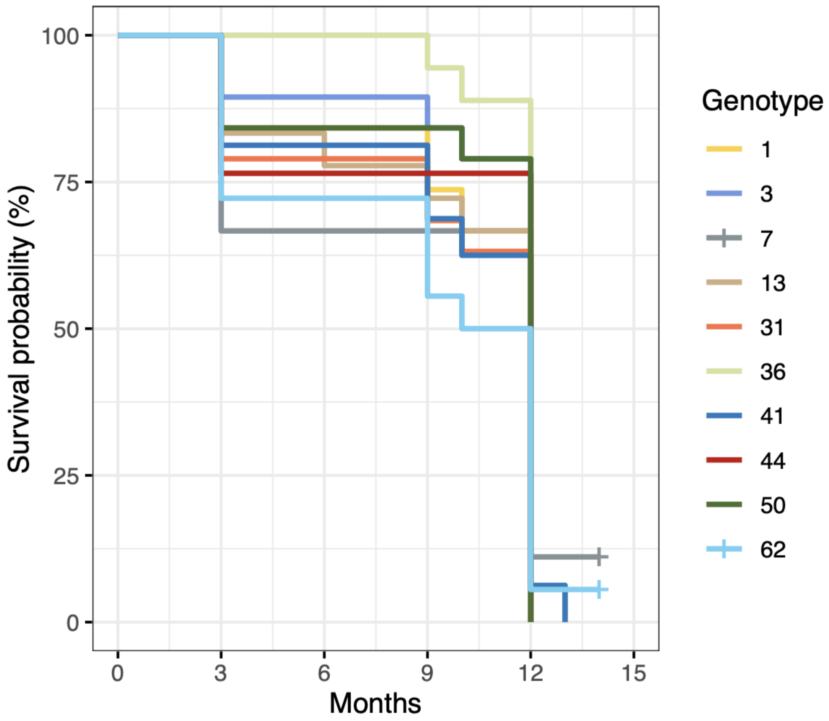


**Figure S9:** Kaplan-Meyer survival curves showing percent survival probability over time for each genotype, represented by color. 9-month mark is the final regular sampling point in June, 12-month mark and onwards represents survival after conclusion of the bleaching event.
